# The minimal endosymbiosis between *Spurilla braziliana* (Aeolidiidae, Nudibranchia) and Symbiodiniaceae

**DOI:** 10.1101/2025.07.17.665303

**Authors:** Hideaki Mizobata, Kenji Tomita, Ryo Yonezawa, Kentaro Hayashi, Kazutoshi Yoshitake, Shigeharu Kinoshita, Shuichi Asakawa

**Affiliations:** Department of Aquatic Bioscience, Graduate School of Agricultural and Life Sciences, the University of Tokyo, Bunkyo-ku, Tokyo, 113-8657 Japan; Technology Advancement Center, Graduate School of Agricultural and Life Sciences, the University of Tokyo, Bunkyo-ku, Tokyo, 113-8657 Japan; College of Bioresource Sciences, Nihon University, Fujisawa, Kanagawa, 252-0880 Japan; School of Marine Biosciences, Kitasato University, Minami-ku, Sagamihara-shi, Kanagawa, 252-0374 Japan

**Keywords:** Endosymbiosis, Nudibranch, Symbiodiniaceae, Photosymbiosis, Histology, Metagenomics

## Abstract

Symbiotic relationships between dinoflagellates of the family Symbiodiniaceae and marine invertebrates underpin the functioning of certain shallow-water marine ecosystems. Although the Symbiodiniaceae-Cladobranchia nudibranch association has been proposed as a promising model for symbiosis research, interspecific variation in the extent of this association remains poorly resolved. Here, we assessed the algal symbiotic characteristics of the nudibranch *Spurilla braziliana*. Histological analyses revealed limited branching of the digestive gland and the presence of intact Symbiodiniaceae cells within lysosome-rich epithelial digestive cells. Metagenomic profiling further showed a complete absence of *Endozoicomonas*—bacteria typically linked to Symbiodiniaceae symbioses—in this species. These findings indicate that *S. braziliana* can harbor Symbiodiniaceae but exhibits only primitive morphological and microbial adaptations to the symbiotic state. Collectively, these results provide new insights into the evolutionary and structural diversity of nudibranch–algal symbioses.

## Introduction

Dinoflagellates of the family Symbiodiniaceae establish symbiotic partnerships with a wide range of marine invertebrates, including cnidarians^1^, molluscs^2^, and foraminiferans^3^. In the most integrated mutualistic associations, hosts provide the algae with inorganic nutrients^1^ and modulate the spectral and angular light field experienced by the symbionts, thereby facilitating photosynthesis^4–6^, while the algae, in turn, often translocate photosynthates that supplement host metabolism^1^. However, in other cases or symbiotic systems, hosts can also digest their symbionts and utilize them as a nutritional resource^7^. Consequently, Symbiodiniaceae–host interactions represent a dynamic continuum ranging from mutualistic to exploitative, depending on environmental and phylogenetic contexts. In the best-studied cnidarian hosts, this association dates back at least 385 million years^8^, and Symbiodiniaceae can, in some cases, supply most of the metabolic substrates required for host energy production^9^. Consequently, Symbiodiniaceae–host symbioses play a foundational role in the functioning of certain shallow-water marine ecosystems, particularly coral reef communities. In recent decades, however, the breakdown of this relationship—most conspicuously manifested as mass bleaching in coral hosts—has become increasingly evident^10^, highlighting the urgent need to clarify the universal mechanisms that govern Symbiodiniaceae–marine invertebrate symbioses.

Cladobranch nudibranchs are considered promising host models for investigating Symbiodiniaceae symbiosis^11^. Their relatively large body size and well-developed anatomical features make them easy to observe, and importantly, the degree of symbiosis varies markedly among species. Some species are capable of maintaining healthy Symbiodiniaceae cells intracellularly for over 200 days^12^, while others retain the algae for only a few weeks or do not internalize them at all^13,14^. Integrating histological information on host traits across closely related taxa with genetic and ecological comparisons offers a promising approach to identify the mechanisms that govern the establishment and maintenance of these associations. Despite this potential, the diversity of algal symbiosis among nudibranchs has been examined in only a limited number of species. Among Cladobranchia, members of the family Aeolidiidae are among the best-studied taxa, and within this family, the model species *Berghia stephanieae* has received particular attention^13–20^. Elucidating the symbiotic state in additional aeolidiid species could therefore fill an important gap in our understanding and establish this lineage as a powerful comparative system for investigating algal symbiosis.In this context, the model genus *Berghia* serves as the primary reference, and within Aeolidiidae, *Spurilla* and *Baeolidia* are among its closest relatives^21^. These three genera form a well-defined clade within the family but differ markedly in their biogeographic distributions^22^: *Berghia* occurs in the Atlantic and Mediterranean^23^, *Baeolidia* is restricted to the Indo-Pacific^22^, and *Spurilla* spans both regions^24^. Since *Berghia* does not inhabit the Pacific Ocean, *Spurilla* offers a valuable opportunity to extend comparative analyses across oceanic regions. Among the five recognized *Spurilla* species, *S. neapolitana* and *S. braziliana* are the most widespread and frequently encountered^24^. The eastern Atlantic species *S. neapolitana* has previously been histologically characterized for its symbiotic association with Symbiodiniaceae^25^, whereas the symbiotic condition of *S. braziliana*, which inhabits both the western Atlantic and the Pacific Ocean, remains poorly understood. In this study, we present the first detailed characterization of the association between *S. braziliana* and Symbiodiniaceae. Through a combination of histological and metagenomic analyses, we demonstrate that this species exhibits a relatively primitive form of symbiosis similar to that observed in other aeolidiid nudibranchs. Our findings thus expand the understanding of Symbiodiniaceae symbiosis within the family and highlight *S. braziliana* as a geographically widespread reference species that can facilitate future comparative studies— particularly in the Pacific region, where established model species are lacking.

## Materials and methods

### Sample collection

Two live specimens of *Spurilla braziliana* (designated as individual A and individual B) were collected from the intertidal zone in Miura, Kanagawa Prefecture, Japan, on June 10 and October 5, 2021, respectively. Individual A is the same specimen previously used in our mitochondrial genome sequencing study^26^ and was preserved in NucleoProtect RNA solution (MACHEREY-NAGEL) immediately after collection. Individual B was maintained in an artificial seawater aquarium (33 g/L, Instant Ocean Premium, NAPQO, Ltd.) equipped with an external filtration system, under constant light conditions and a water temperature of 25°C, for 15 days without feeding. Because this study focused on histological and ultrastructural observations rather than ecological responses, the ambient light intensity and natural habitat conditions were not quantitively measured.

### Fluorescence Microscopy Observation

Fluorescence microscopy was conducted following the same protocol as described in a previous study^27^. The cerata of individual A were excised on the day of sampling (prior to immersion in NucleoProtect RNA) and chemically fixed in 4% paraformaldehyde (FUJIFILM Wako Pure Chemical Corporation). After washing with PBS (FUJIFILM Wako Pure Chemical Corporation), the samples were cleared using the CUBIC Trial Kit (FUJIFILM Wako Pure Chemical Corporation), according to the protocol provided by the manufacturer. During the clearing process, nuclear staining was performed using DAPI solution (DOJINDO LABORATORIES). Chlorophyll autofluorescence from Symbiodiniaceae and DAPI fluorescence were observed under an epifluorescence microscope (KEYENCE BZX-810) equipped with DAPI (EX 360/40 nm, BA 460/50 nm) and TxRed (EX 560/40 nm, BA 630/75 nm) filter cubes (Keyence, Osaka, Japan).

### Light and Transmission Electron Microscopy of Tissue Sections

Transmission electron microscopy (TEM) was carried out following the same protocol as described in a previous study^27^. The cerata, tentacles, and rhinophores of individual B, which had been maintained for 15 days, were dissected and fixed in an aldehyde-based fixative solution [2% paraformaldehyde, 2% glutaraldehyde (FUJIFILM Wako Pure Chemical Corporation), 50 mM HEPES (FUJIFILM Wako Pure Chemical Corporation), 33 g/L Instant Ocean Premium (NAPQO Ltd.)]. After rinsing with distilled water, the tissues were post-fixed in 1% osmium tetroxide (Nisshin EM) at 4°C for 1 hour. Following washing and dehydration through a graded ethanol series, the samples were transferred to propylene oxide and subsequently embedded in epoxy resin (Quetol 812, Nisshin EM). The resin blocks were sectioned using an ultramicrotome (Leica Ultracut UCT). Semi-thin sections (500 nm thick) were stained with toluidine blue (Diagonal GmbH& Co. KG.) and observed under a light microscope (KEYENCE BZX-810). Ultrathin sections (60 nm thick) were stained with 4% aqueous uranyl acetate (Bio-Rad) and Reynolds’ lead citrate (FUJIFILM Wako Pure Chemical Corporation), and observed using a transmission electron microscope (JEOL JEM-1010).

### Identification of the Symbiotic Symbiodiniaceae Genus

Raw whole-genome shotgun sequence reads from individual A (GenBank Accession ID: DRR450785)^26^ were assembled using CLC Genomics Workbench version 8.5 (QIAGEN Aarhus A/S). To identify symbiotic Symbiodiniaceae sequences, we used the reference dataset published by Pochon et al.^28^ and conducted blastn searches against all contigs in the primary assembly for six genetic markers: nuclear *28S rRNA*, nuclear *elf2*, mitochondrial *COI*, mitochondrial *cob*, chloroplast *23S rRNA*, and chloroplast *psbA* genes. Contigs identified as Symbiodiniaceae-derived were subsequently aligned to the reference dataset using MAFFT v7.490^29,30^ on Geneious Prime. Based on these alignments, maximum likelihood phylogenetic trees were constructed using IQ-TREE version 2.3.3^31^ with 1000 bootstrap replicates. To reduce overestimation of branch support, the -bnni option was applied. Additionally, SH-aLRT^32^ and aBayes^33^ tests were performed to provide a more stringent assessment of node support. The final phylogenetic trees were visualized using Interactive Tree of Life version 6^34^.

### Visualization of the Hologenome Structure

Raw reads were mapped to the primary assembly using BWA-MEM^35^, and the resulting alignments were processed into BAM files with samtools version 1.17^36^. The BAM files were analyzed with CoverM version 0.6.1 (https://github.com/wwood/CoverM) to calculate the mean coverage for each contig. Contig lengths were determined using seqkit version 2.3.0^37^ and GNU Awk version 4.0.2 (https://www.gnu.org/software/gawk/). The 4-mer composition of each contig was calculated using Perl scripts included in the Multi-metagenome^38^. Feature data from each contig were visualized as scatter plots using the matplotlib package in Python version 3.10.4. Contig coverage and length were used as the x- and y-axis coordinates, respectively. The 256-dimensional 4-mer vectors were reduced to three dimensions using UMAP (umap-learn module, version 0.5.3)^39^, normalized, and converted into RGB color values that were applied to the data points in the plot.

### Identification of Organisms Present in the Holobiont

Each contig in the primary assembly was taxonomically characterized by homology search against the NCBI nr database (downloaded on May 19, 2021; https://ftp.ncbi.nlm.nih.gov/blast/db/) using DIAMOND version 2.0.9^40^. Only hits with an E-value of 0.0 and a bit score within 95% of the top hit were retained. Taxonomic assignments for each contig were determined using the lowest common ancestor (LCA) approach. A scatter plot was then redrawn using contig coverage and length as the x- and y-axes, respectively, with each contig colored according to its assigned taxon.

Furthermore, we performed a homology search using blastn against the NCBI nt database with the raw reads from WGS. A total of 200,000 reads (100,000 reads each from R1 and R2) with any hits were extracted, and from these, reads whose top hit had an e-value < 1e-50 were further selected. For the extracted reads, hits with bit scores ≥95% of the top hit were considered true hits.

## Results

### Whole-Mount Localization of Symbiodiniaceae in the Cerata

Whole-mount fluorescence microscopy revealed the internal structure of the ceras in *Spurilla braziliana*. While most cells within the ceras exhibited only blue fluorescence from DAPI staining, red autofluorescence—potentially derived from chlorophyll of Symbiodiniaceae—was observed in the slightly branched digestive gland located at the center of the ceras (Fig. 1). In contrast, no autofluorescence was detected in the cnidosacs, which are extensions of the digestive gland.

**Figure 1.**
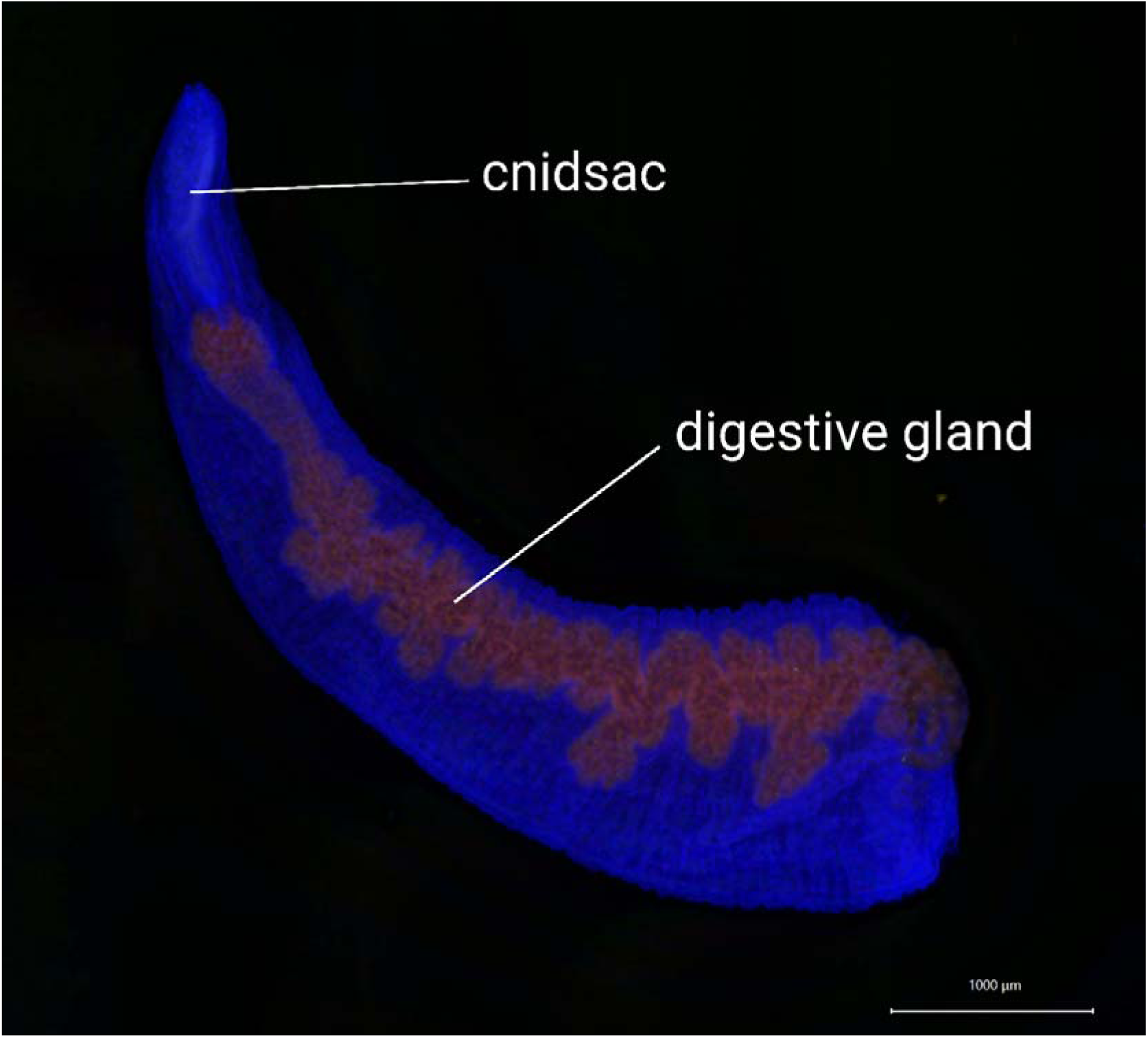
Whole-mount fluorescence microscopy image of a *Spurilla braziliana* ceras. Red autofluorescence is visible exclusively in the digestive gland. Blue fluorescence indicates DAPI staining. The exposure times were 1/8.5 s for the green and red channels, 1/80 s for the blue channel.

### Light and Transmission Electron Microscopy of Tissue Sections

Observation of toluidine blue-stained sections under a light microscope allowed examination of the two-dimensional tissue architecture in the cerata, tentacles, and rhinophores of *S. braziliana* (Fig. 2). A gently branched digestive gland extended through the center of each ceras, and multiple Symbiodiniaceae cells were observed in its epithelial layer (Fig. 2a). All five branches of the digestive gland exhibited similar structure, although algal cells were not observed in the central branch within the limits of our observations. No algal cells were detected outside the digestive gland epithelium within the ceras. In both the tentacles and rhinophores, no penetration of the digestive gland was observed, and no algal cells were present (Fig. 2b, c).

**Figure 2.**
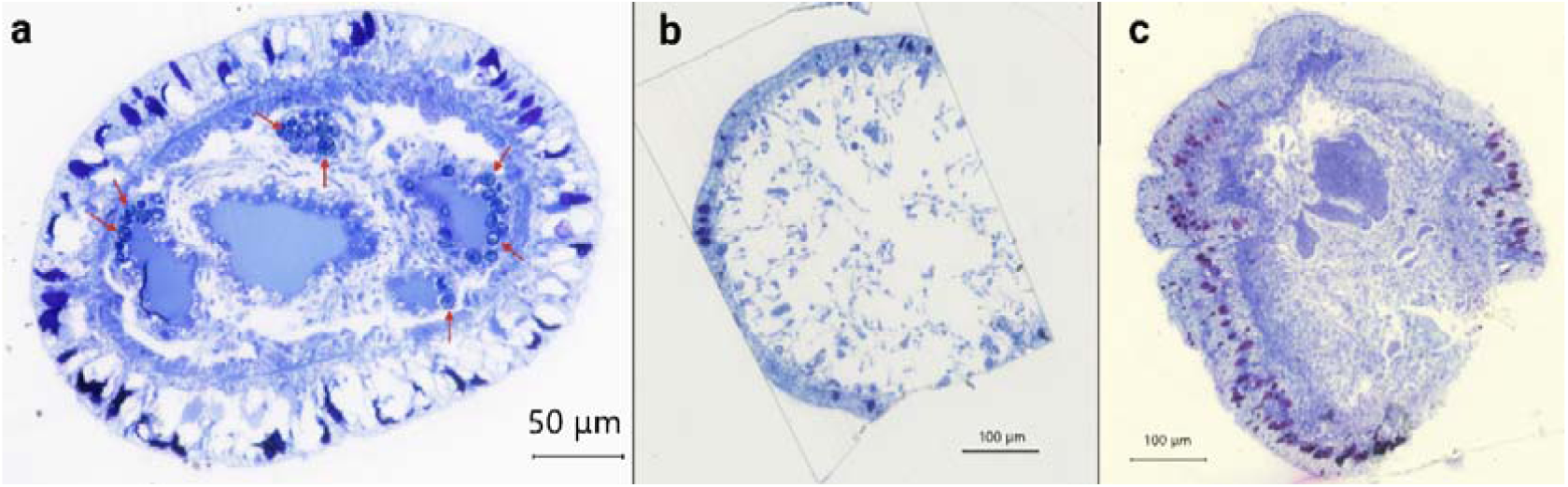
Toluidine blue-stained sections of various tissues in *Spurilla braziliana*. a. Cross-section of a ceras. The digestive gland divides into five diverticula, within which algal cells are located in the epithelium. Arrowheads indicate algal cells. b. Cross-section of a tentacle. No penetration of the digestive gland or presence of algal cells is observed. c. Cross-section of a rhinophore. No penetration of the digestive gland or presence of algal cells is observed.

Transmission electron microscopy (TEM) was used to examine the ultrastructure of algal symbiosis within the digestive gland epithelium of the ceras in *S. braziliana* (Fig. 3). The lumen of the digestive gland was lined with cilia or microvilli, and its epithelial cells contained numerous lysosomes and Symbiodiniaceae cells (Fig. 3a). Many of the algal cells appeared to retain intact cellular architecture, with clearly distinguishable internal structures, such as nucleus, pyrenoids, chloroplasts, lipid droplets, and starches (Fig. 3c). Some of the algal cells were in the stage of cell division^41^ (Fig. 3d). Only a few degraded algal cells were observed.

**Figure 3.**
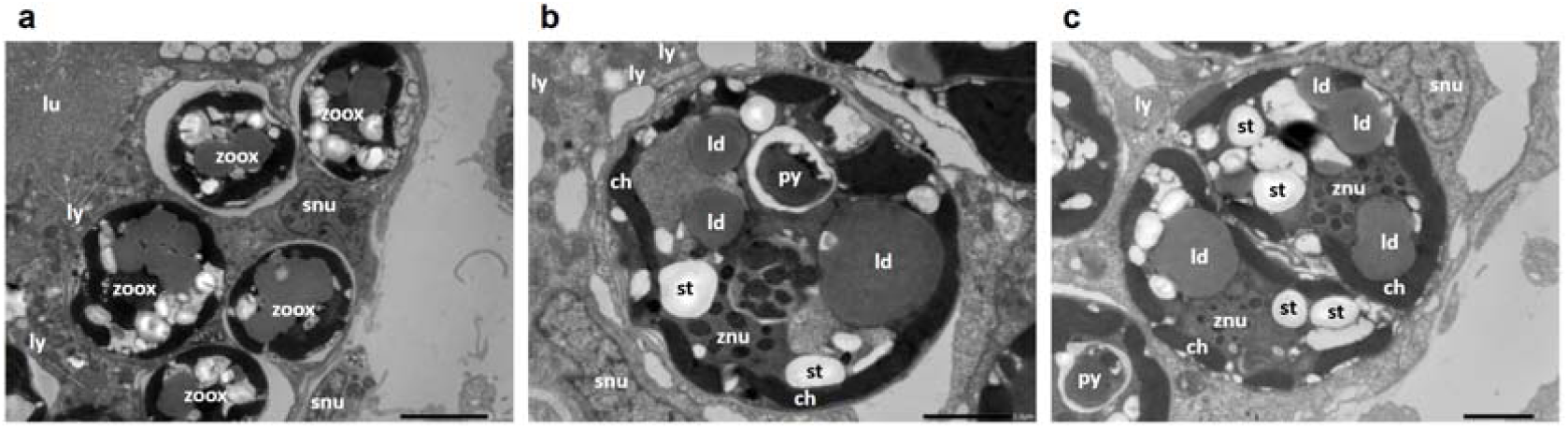
TEM images of the digestive gland within a ceras. a. Algal cells and lysosomes within the epithelial cells. Scale bar: 5 µm. b. Cellular structure of a single Symbiodiniaceae cell within a sea slug cell. Scale bar: 2 µm. c. Symbiodiniaceae cells in the process of cell division. Scale bar: 2 µm. ch: chloroplast, ld: lipid droplet, lu: digestive gland lumen, ly: lysosome, py: pyrenoid, snu: nuclei of sea slug cells, st: starch granules, znu: nuclei of Symbiodiniaceae, zoox: Symbiodiniaceae cell

### Clade Affiliation of Symbiodiniaceae in *S. braziliana*

Using blastn searches followed by manual alignment inspection, one contig each was identified as homologous to the mitochondrial *COI* and *cob* genes based on similarity to reference sequences. In contrast, the remaining four genes (nuclear and chloroplast markers) either produced no significant hits or showed clearly low sequence similarity. Phylogenetic trees were constructed for the two contigs with confirmed homology to the reference dataset (Fig. 4). In both the *COI* and *cob* gene trees, the contigs derived from *S. braziliana* formed a monophyletic group with *Symbiodinium* (clade A) species, indicating that the symbiotic algae associated with *S. braziliana* belong to *Symbiodinium*.

**Figure 4.**
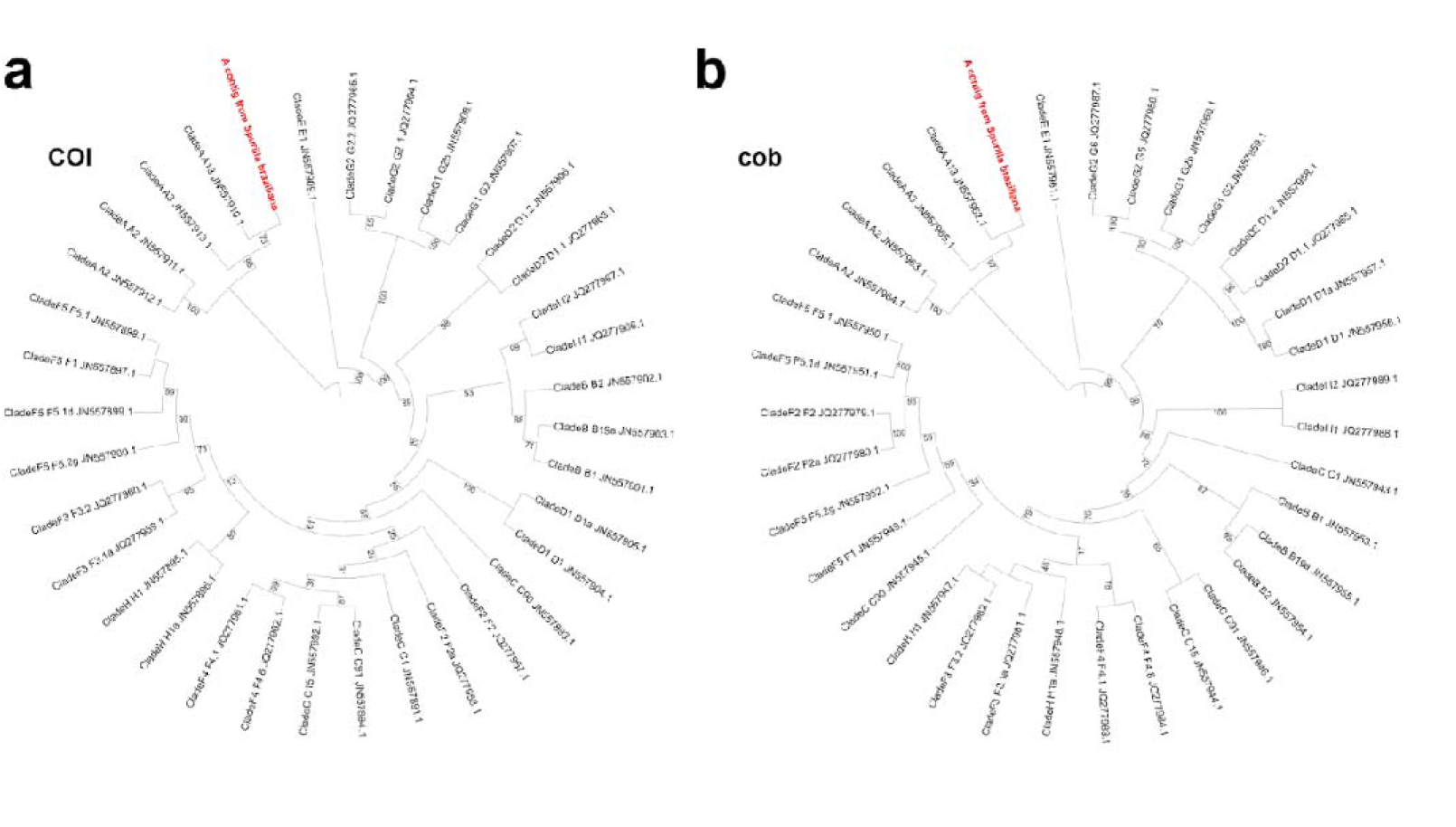
Phylogenetic placement of Symbiodiniaceae-derived sequences detected in *Spurilla braziliana*. Branch lengths are not to scale for clarity. For full trees with branch lengths, see Supplementary Data 1. a. *COI* gene; b. *cob* gene.

### Hologenomic Structure in *Spurilla braziliana*

In the *S. braziliana*–Symbiodiniaceae symbiotic system, not only the dinoflagellates but also a broader microbiome may contribute to symbiotic adaptation. To explore this possibility, we visualized the major microbiome components present within the nudibranch using previously obtained whole-genome shotgun sequencing data (Fig. 5). Each contig from the primary assembly was plotted using its coverage and length as spatial coordinates, and its color was determined by dimensional reduction (3D UMAP) of its 4-mer sequence composition. This visualization revealed four clearly distinct peaks, each representing contigs with differing compositional properties (Fig. 5a).

Using DIAMOND blastx searches and taxonomic assignment via the lowest common ancestor (LCA) method, we were able to infer the likely source organisms for 1,294 out of 850,960 contigs in the primary assembly. Among these, taxa represented by more than 50 contigs included *Vibrio* (977 contigs), *Photobacterium* (134 contigs), and Gastropoda (77 contigs), corresponding to the host itself. An additional 56 bacterial contigs were attributed to taxa other than the two dominant genera. Other notable assignments included Embryophyta (24 contigs) and Symbiodiniaceae (4 contigs). When taxonomic assignments were overlaid onto the coverage–contig length scatter plot, it became evident that distinct taxa underlie the four prominent peaks observed in Fig. 5a (Fig. 5b). Peak 1 was primarily composed of contigs assigned to the genus *Photobacterium*, while Peak 2 was dominated by *Vibrio*. Peak 3 corresponded to Gastropoda, reflecting sequences derived from the host organism. Peak 4, though less sharply defined, exhibited the highest coverage and was attributed to a heterogeneous group of other bacterial taxa. Contigs matching Symbiodiniaceae or other non-bacterial organisms did not form distinct peaks in the plot.

Among the bacteria other than *Photobacterium* and *Vibrio*, seven families were identified through LCA-based analysis at the family level: Cyclobacteriaceae, Enterobacteriaceae, Flammeovirgaceae, Marinifilaceae, Marinilabiliaceae, Prolixibacteraceae, and Pseudomonadaceae. Of these, contigs assigned to Enterobacteriaceae and Pseudomonadaceae tended to have relatively low coverage (Fig. 5c).

On the other hand, *Endozoicomonas*, which is of particular interest in the context of endosymbiosis, was not detected. To confirm the absence of *Endozoicomonas*, we performed a blastn search using the raw reads from WGS. As a result, among the subset of raw reads (200,000 reads) that produced any hits, no reads showed high-scoring hits to Endozoicomonadaceae, including *Endozoicomonas*.

**Figure 5.**
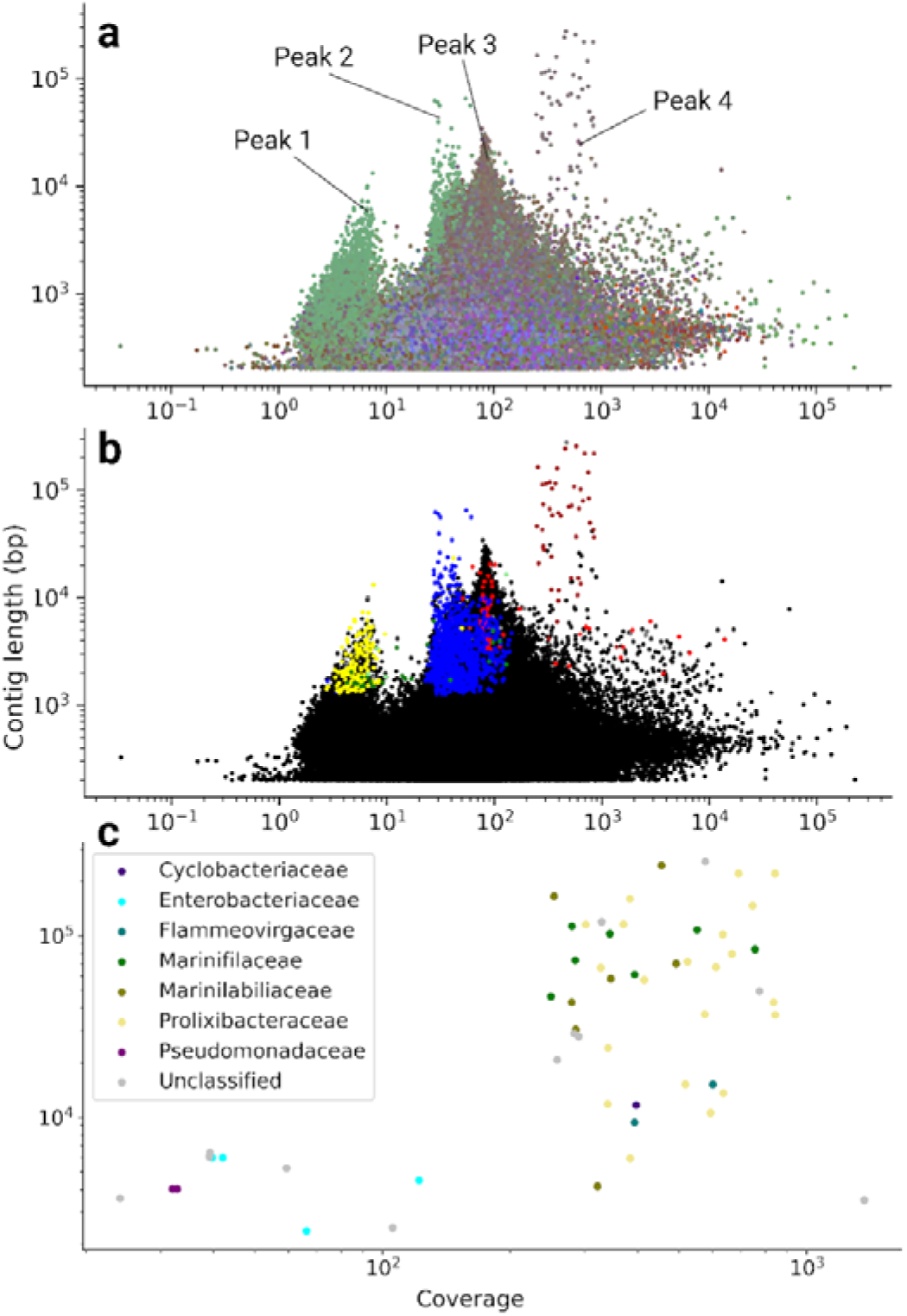
Hologenomic structure of the *Spurilla braziliana* – Symbiodiniaceae holobiont. a. Coverage–contig length scatter plot colored by sequence composition. b. Coverage–contig length scatter plot colored by taxonomic assignment. Each peak corresponds to a different biological origin. Black: no hit; yellow: *Photobacterium*; blue: *Vibrio*; red: Gastropoda; brown: other bacteria; green: Embryophyta; light green: Symbiodiniaceae; gray: others. c. Coverage–contig length scatter plot focusing on contigs assigned to bacterial taxa.

## Discussion

### Symbiotic Structure of Spurilla *braziliana*

In Cladobranchia, symbiotic associations with Symbiodiniaceae have been reported in four superfamilies: Aeolidioidea, Fionoidea, Arminoidea, and Dendronotoidea. The ability to establish such symbioses is thought to have evolved independently multiple times. Generally, algal symbionts reside intracellularly within the epithelial cells of the nudibranch’s digestive gland^11^.

A highly branched digestive gland extending into the cerata is a defining feature shared by species capable of medium-to long-term retention of Symbiodiniaceae. Representative examples include *Pteraeolidia semperi* and *Phyllodesmium briareum* (Aeolidioidea) as well as *Melibe engeli* and *Melibe pilosa* (Dendronotoidea). In these species, the ceratal digestive gland system consists of extensively branched “fine tubules” that contain specialized epithelial “carrier cells”, which are structurally distinct from those of the main digestive gland^12,27,42^. It should be noted, however, that certain long-term symbiotic species, such as *Phyllodesmium longicirrum*, lack fine tubules despite their capacity for prolonged algal retention^12^. Detailed ultrastructural observations of fine tubules are currently available for three species: *Pteraeolidia semperi*, *Melibe pilosa*, and *Melibe sp.* In *Pteraeolidia semperi*, the epithelial cells of the main digestive gland (the “central duct”) generally lack Symbiodiniaceae and contain abundant lysosomes, whereas the carrier cells forming the fine tubules contain the algae but few lysosomes^27^. Also in *Melibe*, the carrier cells lack lysosomes, although detailed images of the central duct have not been presented^42^. These features were not described in the original studies but are clearly discernible from the available TEM images. Epithelial cells rich in lysosomes are widely recognized throughout Mollusca as typical digestive cells^43,44^, suggesting that, in these taxa, the digestive gland epithelium has functionally differentiated into general digestive cells and carrier cells specialized for endosymbiosis. This differentiation represents an evolutionarily advanced and symbiotically adaptive condition. In most other cladobranch species, however, the ultrastructural organization of the “fine tubules” has not yet been examined in detail. In these species, it remains unclear whether the descriptions “fine tubules” simply represent slender extensions of the digestive gland composed of ordinary digestive cells, or whether they constitute specialized organs formed by carrier cells adapted for symbiosis.

Within the family Aeolidiidae (Aeolidioidea) which includes *Spurilla braziliana*^21–24,45–50^, symbiotic associations with Symbiodiniaceae have been documented in 16 species, whereas 2 species are reported to lack such symbionts^13,23–25,42,48,51–56^ (Fig. 6). Among the species in which the epithelial organization of the digestive gland has been examined, namely *Aeolidiopsis ransoni*^52^, *Aeolidiopsis harrietae*^52^, *Baeolidia australis*^52^, *Baeolidia moebii*^42,52^, *Berghia verrucicornis*^51^, *Bulbaeolidia alba*^48,57^, and *Spurilla neapolitana*^25^, no fine tubule-like structures have been observed within the cerata. In these nudibranchs, Symbiodiniaceae occur within the epithelial cells of relatively large branches of the digestive gland. Although the ultrastructure of these epithelial cells has not been extensively characterized, available micrographs for *Baeolidia moebii*^42^, *Berghia verrucicornis*^51^, *Berghia stephanieae*^11,18^, and *Spurilla neapolitana*^25^ reveal lysosome-like structures within the algae-retaining cells. This indicates that, in these species, the digestive cells themselves serve as hosts for Symbiodiniaceae, representing an unspecialized form of symbiosis in which carrier cells have not yet differentiated.

**Figure 6.**
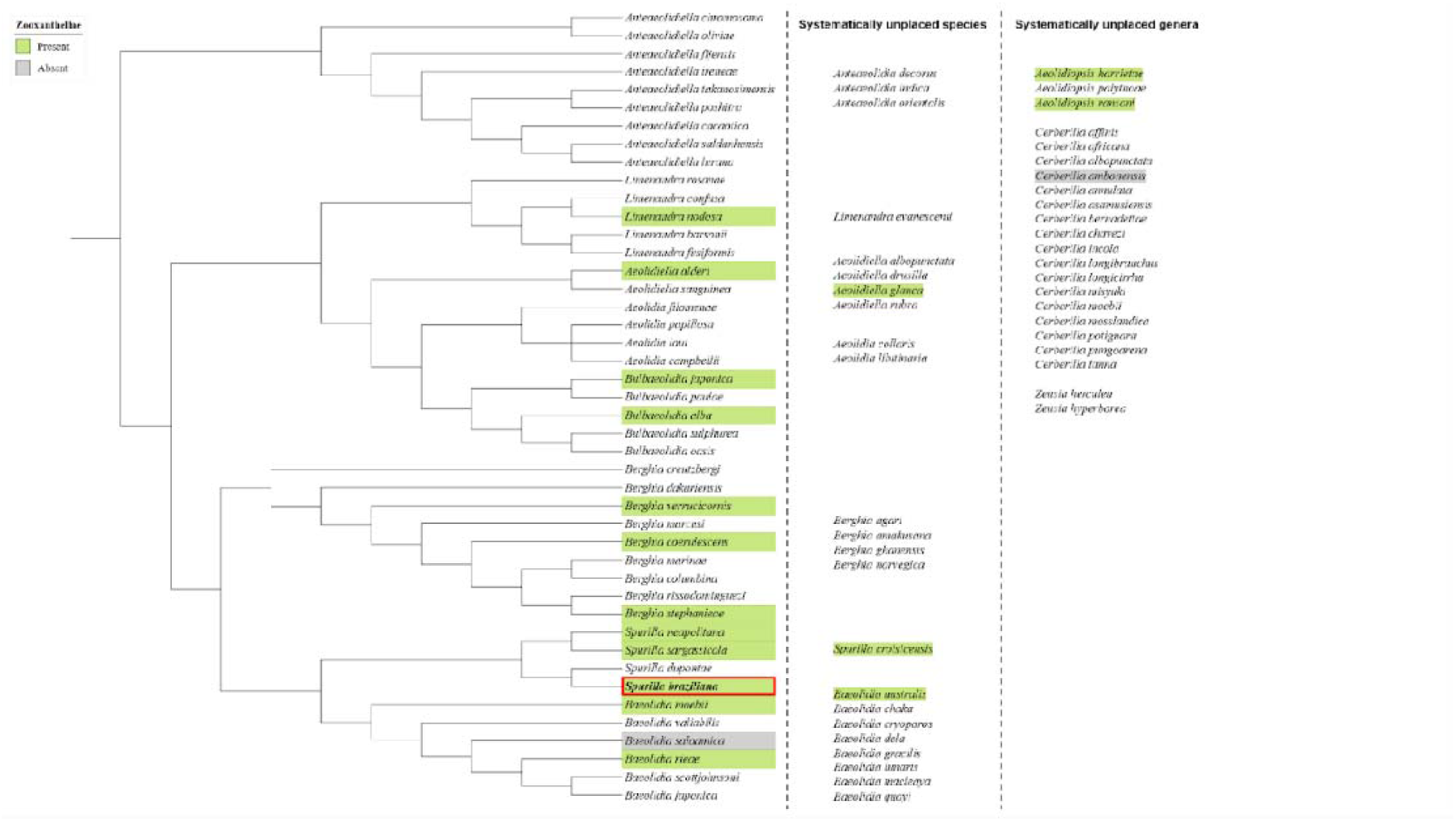
Presence of Symbiodiniaceae in the family Aeolidiidae. The species list of Aeolidiidae follows MolluscaBase as accessed via WoRMS on June 30, 2025^50^. Green: species for which the presence of Symbiodiniaceae has been reported. Gray: species for which the absence of Symbiodiniaceae has been reported. No color: species for which neither presence nor absence has been confirmed. Red box: present study. See Supplementary Table 1 for references supporting phylogenetic placement and symbiont status.

In *Spurilla braziliana*, the digestive gland within the cerata was limited in its branching (Fig. 1), and no discernible structural variation was detected among the five branches examined (Fig. 2a). Symbiodiniaceae were retained within lysosome-rich digestive cells (Fig. 3a), consistent with the absence of epithelial cell differentiation. These findings indicate that the morphology of the symbiotic cells in this species represents an unspecialized form consistent with that typically observed in the family Aeolidiidae. However, *S. braziliana* also exhibits a distinct feature within the family. In other aeolidiid nudibranchs known to harbor algal symbionts, namely *Aeolidiopsis ransoni*^52^, *Aeolidiopsis harrietae*^52^, *Baeolidia australis*^52^, *Baeolidia moebii*^52^, and *Spurilla neapolitana*^25^, the digestive gland has been reported to extend into the rhinophores and/or oral tentacles. In contrast, such extensions were not observed in *S. braziliana* (Fig. 2b, c), suggesting that this species possesses a particularly low degree of digestive gland branching compared with its close relatives.

Within the superfamily Aeolidioidea, and particularly within the family Aeolidiidae, a correlation has been proposed between the degree of digestive gland branching and the ability to retain symbiotic algae^12,56,58^. The relatively simple and undifferentiated epithelial organization of the digestive gland observed in *Spurilla braziliana*, which appears comparable to or even less complex than that of its close relatives, could therefore indicate that this species has an algal-retention capacity similar to that of other aeolidiid nudibranchs. Members of the Aeolidiidae typically maintain their algal symbionts for short to intermediate periods. Short-term retention has been reported in *Berghia stephanieae*^13,14^ (3–8 days) and *Aeolidia papillosa*^55^ (11 days), whereas intermediate-term retention has been documented in *Aeolidiella alba*^56^ (40 days), *Baeolidia australis*^56^ (≥22 days), and *Limenandra nodosa*^56^ (≥22 days). In the present study, numerous Symbiodiniaceae cells with well-preserved intracellular structures, most notably intact lipid droplets and starch granules, were still present within the host cells of *S. braziliana* after 15 days of starvation (Figs. 3a, b). Moreover, some algal cells were observed undergoing division (Fig. 3c). These observations suggest that *S. braziliana* provides a relatively favorable intracellular environment for Symbiodiniaceae survival and that this species likely exhibits at least a short-to intermediate-term algal retention capability comparable to that of other aeolidiid nudibranchs.

During our field surveys, *Pteraeolidia semperi* was also recorded in the same coastal area. Whereas *Pteraeolidia semperi* individuals were collected from sunlit habitats, *Spurilla braziliana* was found beneath rocks. This negative phototactic behavior has also been reported in certain other nudibranchs harboring Symbiodiniaceae, such as *Berghia stephanieae*^59^ and *Phidiana lynceus*^60^. As described previously, *Berghia stephanieae* retains algal symbionts for only about one week^13,14^, whereas *Phidiana lynceus* can maintain them for a longer period but dies within approximately 17 days without obtaining metabolic benefits from the symbionts^60^. The negative phototaxis observed in *S. braziliana* therefore suggests that, like these species, it may not be ecologically adapted to photosymbiosis.

However, light conditions, morphology, and algal abundance were neither standardized nor quantitatively assessed in the present study, precluding any definitive conclusions regarding the algal retention capacity or the degree of symbiotic adaptation in *Spurilla braziliana*. Furthermore, the present observations were based on only two biological specimens. Given that multiple phenotypes have been reported within this species^24^, individuals differing in phenotype may exhibit variation in symbiotic capacity. Future studies should secure a larger number of specimens and perform a detailed characterization of phenotypic variation. Quantitative evaluation of the symbiosis should involve both metabolic profiling and assessment of algal retention ability using pulse amplification modulation (PAM) fluorometry, integrated with morphological indicators such as the branching pattern of the digestive gland.

In this section, we summarized the morphological features and algal-retention capacities reported for aeolidiid nudibranchs (Fig. 6) and characterized the symbiotic morphology of *Spurilla braziliana* in detail. Within Cladobranchia, algal symbiosis is thought to have evolved multiple times independently^11^. Therefore, assessments at relatively shallow phylogenetic levels, such as within a family, are better suited for discussing how a single ancestral symbiotic state may have diversified. Because both symbiotic and non-symbiotic species are represented and histological information is relatively comprehensive, Aeolidiidae provide a coherent model system for comparative and evolutionary studies of algal symbiosis. Among aeolidiids, *S. braziliana* is characterized by the absence of digestive-gland extensions into the rhinophores and tentacles. Although this feature may not be functionally significant, it provides a convenient reference point for interspecific transcriptomic comparisons of the rhinophores or tentacles aimed at identifying genes associated with algal symbiosis. In addition, *Baeolidia salaamica*, a non-symbiotic species that occurs within a clade otherwise composed of symbiotic taxa (Fig. 6), may represents another valuable comparative species for clarifying how the loss of symbiosis occurred within Aeolidiidae^52^.

### Microbiome Composition in *Spurilla braziliana*

From the whole-genome shotgun (WGS) assembly of *S. braziliana*, single contigs corresponding to the *COI* and *cob* genes of dinoflagellates were identified. Phylogenetic analysis showed that both sequences belong to the genus *Symbiodinium* (clade A) (Fig. 4). Nudibranchs within the suborder Cladobranchia have been reported to form symbioses with Symbiodiniaceae clades A through D— *Symbiodinium*, *Breviolum*, *Cladocopium*, and *Durusdinium*—with the specific clade often reflecting the host’s geographic distribution^61–67^. *Symbiodinium* (clade A) is frequently detected in nudibranchs from temperate regions^62^ and has previously been reported in *Pteraeolidia semperi* from Hayama^61^, a locality very close to the present sampling site. This finding aligns with known biogeographic trends in Cladobranchia–Symbiodiniaceae associations.

Regarding the bacterial community of *S. braziliana*, contigs assigned to the genera *Photobacterium* and *Vibrio* were detected at relatively low coverage. These genera are commonly reported as intestinal bacteria in marine gastropods^68^. In addition, members of the families Cyclobacteriaceae, Enterobacteriaceae, Flammeovirgaceae, Marinifilaceae, Marinilabiliaceae, Prolixibacteraceae, and Pseudomonadaceae were identified. Among these, Enterobacteriaceae and Pseudomonadaceae have been reported to co-occur with certain clades of Symbiodiniaceae in coral hosts^69^. However, their functional roles remain poorly understood, and the study has specifically noted a lack of co-occurrence with *Symbiodinium* (clade A)^69^. Therefore, caution is warranted in interpreting any direct contribution of these bacterial taxa to algal symbiosis in *S. braziliana*.

Among bacterial taxa associated with photosymbiosis, the genus *Endozoicomonas* has attracted the most attention. In cnidarian hosts, members of this family are thought to positively influence algal symbiosis by modulating host immunity and supplying essential metabolites that support Symbiodiniaceae photosynthesis^70–72^. In nudibranch hosts as well, *Endozoicomonas* has been identified in species with both advanced (*Pteraeolidia semperi*)^73^ and rudimentary (*Berghia stephanieae*)^19^ symbiosis. However, in the present study, no contigs assigned to Endozoicomonadaceae were detected from *S. braziliana*. This absence of symbiosis-associated bacterial taxa may further support the interpretation that *S. braziliana* possesses a low level of symbiotic adaptation. That said, the actual functional roles of bacterial communities in marine invertebrate–Symbiodiniaceae symbioses remain poorly understood. As our understanding of host– symbiont–microbiome interactions deepens, it will become possible to more precisely evaluate the symbiotic potential of species like *S. braziliana*.

## Conclusion

The study of nudibranch-Symbiodiniaceae symbiosis remains in its early stages, and the diversity of symbiotic capacity across species is still poorly understood. In this study, we conducted the first histological and metagenomic analyses of endosymbiosis in *Spurilla braziliana*, a species previously unexamined in this context. The combination of a simple digestive morphology for algal retention and the absence of bacterial taxa typically associated with algal symbiosis suggests that *S. braziliana* exhibits primitive algal symbiotic capacity. Given its broad distribution encompassing both the Atlantic and Pacific Oceans, *S. braziliana* further offers practical value for expanding comparative research into regions where established model species such as *Berghia* are absent. Collectively, these insights refine our understanding of the evolutionary and structural diversity of Symbiodiniaceae symbioses within Aeolidiidae. As comparative analyses of related species continue to expand, we will be better equipped to bridge interspecific and intergeneric gaps and identify the key factors underlying the emergence of symbiotic capacity in marine invertebrates.

## Author contributions

H.M. conceived the study. H.M., R.Y., and K.H. conducted sampling and maintenance of live nudibranchs. H.M., K.T., and K.H. performed histological observations. H.M. carried out microbiome analyses. H.M. interpreted the results and wrote the manuscript. All authors contributed to discussions and revised the manuscript.

## Competeing Interests

The authors declare that they have no competing interests.

## Reference

1. Jacobovitz, M. R., Hambleton, E. A. & Guse, A. Unlocking the Complex Cell Biology of Coral–Dinoflagellate Symbiosis: A Model Systems Approach. Annu. Rev. Genet. 57, 411–434 (2023).

2. Mies, M. Evolution, diversity, distribution and the endangered future of the giant clam–Symbiodiniaceae association. Coral Reefs 38, 1067–1084 (2019).

3. LeKieffre, C. et al. Assimilation, translocation, and utilization of carbon between photosynthetic symbiotic dinoflagellates and their planktic foraminifera host. Mar. Biol. 165, 104 (2018).

4. Wangpraseurt, D., Larkum, A. W. D., Ralph, P. J. & Kühl, M. Light gradients and optical microniches in coral tissues. Front. Microbiol. 3, (2012).

5. Rossbach, S., Subedi, R. C., Ng, T. K., Ooi, B. S. & Duarte, C. M. Iridocytes Mediate Photonic Cooperation Between Giant Clams (Tridacninae) and Their Photosynthetic Symbionts. Front. Mar. Sci. 7, 465 (2020).

6. Bollati, E. et al. Green fluorescent protein-like pigments optimise the internal light environment in symbiotic reef-building corals. eLife 11, e73521 (2022).

7. Wiedenmann, J. et al. Reef-building corals farm and feed on their photosynthetic symbionts. Nature 620, 1018–1024 (2023).

8. Jung, J. et al. Coral photosymbiosis on Mid-Devonian reefs. Nature 636, 647–653 (2024).

9. Grottoli, A. G., Rodrigues, L. J. & Palardy, J. E. Heterotrophic plasticity and resilience in bleached corals. Nature 440, 1186–1189 (2006).

10. Hughes, T. P. et al. Global warming and recurrent mass bleaching of corals. Nature 543, 373–377 (2017).

11. Rola, M. et al. Cladobranchia (Gastropoda, Nudibranchia) as a Promising Model to Understand the Molecular Evolution of Photosymbiosis in Animals. Front. Mar. Sci. 8, 745644 (2022).

12. Burghardt, I. Symbiosis between Symbiodinium (Dinophyceae) and various taxa of Nudibranchia (Mollusca: Gastropoda), with analyses of long-term retention. Org. Divers. Evol. 8, 66–76 (2008).

13. Monteiro, E. A., Güth, A. Z., Banha, T. N. S., Sumida, P. Y. G. & Mies, M. Evidence against mutualism in an aeolid nudibranch associated with Symbiodiniaceae dinoflagellates. Symbiosis 79, 183–189 (2019).

14. Silva, R. X. G., Cartaxana, P. & Calado, R. Prevalence and Photobiology of Photosynthetic Dinoflagellate Endosymbionts in the Nudibranch *Berghia stephanieae*. Animals 11, 2200 (2021).

15. Mies, M. et al. Expression of a symbiosis-specific gene in *Symbiodinium* type A1 associated with coral, nudibranch and giant clam larvae. R. Soc. Open Sci. 4, 170253 (2017).

16. Clavijo, J. M. et al. The nudibranch *Berghia stephanieae* (Valdés, 2005) is not able to initiate a functional symbiosome-like environment to maintain *Breviolum minutum* (J.E.Parkinson & LaJeunesse 2018). Front. Mar. Sci. 9, 934307 (2022).

17. Silva, R. X. G., Madeira, D., Cartaxana, P. & Calado, R. Assessing the Trophic Impact of Bleaching: The Model Pair *Berghia stephanieae*/*Exaiptasia diaphana*. Animals 13, 291 (2023).

18. Borgstein, N. M., Van Der Meij, S. E. T., Christa, G. & Laetz, E. M. J. Potential energetic and oxygenic benefits to unstable photosymbiosis in the cladobranch slug, *Berghia stephanieae* (Nudibranchia, Aeolidiidae). Mar. Biol. Res. 20, 45–58 (2024).

19. Sickinger, C. et al. Microbiome origin and stress-related changes in bacterial abundance of the photosymbiotic sea slug Berghia stephanieae (Á. Valdés, 2005). Symbiosis 93, 177–192 (2024).

20. Goodheart, J. A. et al. A chromosome-level genome for the nudibranch gastropod *Berghia stephanieae* helps parse clade-specific gene expression in novel and conserved phenotypes. BMC Biol. 22, 9 (2024).

21. Carmona, L., Pola, M., Gosliner, T. M. & Cervera, J. L. A Tale That Morphology Fails to Tell: A Molecular Phylogeny of Aeolidiidae (Aeolidida, Nudibranchia, Gastropoda). PLoS ONE 8, e63000 (2013).

22. Carmona, L., Pola, M., Gosliner, T. M. & Cervera, J. L. Review of *Baeolidia*, the largest genus of Aeolidiidae (Mollusca: Nudibranchia), with the description of five new species. Zootaxa 3802, (2014).

23. Carmona, L., Pola, M., Gosliner, T. M. & Cervera, J. L. The Atlantic-Mediterranean genus Berghia Trinchese, 1877 (Nudibranchia: Aeolidiidae): taxonomic review and phylogenetic analysis. J. Molluscan Stud. 80, 482–498 (2014).

24. Carmona, L. et al. Untangling the Spurilla neapolitana (Delle Chiaje, 1841) species complex: a review of the genus Spurilla Bergh, 1864 (Mollusca: Nudibranchia: Aeolidiidae). Zool. J. Linn. Soc. https://doi.org/10.1111/zoj12098 (2014) doi:10.1111/zoj12098.

25. Marin, A. & Ros, J. Presence of Intracellular Zooxanthellae in Mediterranean Nudibranchs. J. Molluscan Stud. 57, 87–101 (1991).

26. Mizobata, H. et al. The complete mitochondrial genome of *Spurilla braziliana* MacFarland 1909 (Nudibranchia, Aeolidiidae). Mitochondrial DNA Part B 8, 862– 866 (2023).

27. Mizobata, H. et al. The highly developed symbiotic system between the solar-powered nudibranch *Pteraeolidia semperi* and Symbiodiniacean algae. iScience 26, 108464 (2023).

28. Pochon, X., Putnam, H. M. & Gates, R. D. Multi-gene analysis of *Symbiodinium* dinoflagellates: a perspective on rarity, symbiosis, and evolution. PeerJ 2, e394 (2014).

29. Katoh, K. MAFFT: a novel method for rapid multiple sequence alignment based on fast Fourier transform. Nucleic Acids Res. 30, 3059–3066 (2002).

30. Katoh, K. & Standley, D. M. MAFFT Multiple Sequence Alignment Software Version 7: Improvements in Performance and Usability. Mol. Biol. Evol. 30, 772–780 (2013).

31. Minh, B. Q. et al. IQ-TREE 2: New Models and Efficient Methods for Phylogenetic Inference in the Genomic Era. Mol. Biol. Evol. 37, 1530–1534 (2020).

32. Guindon, S. et al. New Algorithms and Methods to Estimate Maximum-Likelihood Phylogenies: Assessing the Performance of PhyML 3.0. Syst. Biol. 59, 307–321 (2010).

33. Anisimova, M. et al. Survey of Branch Support Methods Demonstrates Accuracy, Power, and Robustness of Fast Likelihood-based Approximation Schemes. Syst. Biol. 60, 685–699 (2011).

34. Letunic, I. & Bork, P. Interactive Tree Of Life (iTOL) v5: an online tool for phylogenetic tree display and annotation. Nucleic Acids Res. 49, W293–W296 (2021).

35. Li, H. Aligning sequence reads, clone sequences and assembly contigs with BWA-MEM. ArXiv Genomics https://doi.org/10.6084/m9.figshare.963153.v1 (2013) doi:10.6084/m9.figshare.963153.v1.

36. Danecek, P. et al. Twelve years of SAMtools and BCFtools. GigaScience 10, giab008 (2021).

37. Shen, W., Le, S., Li, Y. & Hu, F. SeqKit: A Cross-Platform and Ultrafast Toolkit for FASTA/Q File Manipulation. PLOS ONE 11, e0163962 (2016).

38. Albertsen, M. et al. Genome sequences of rare, uncultured bacteria obtained by differential coverage binning of multiple metagenomes. Nat. Biotechnol. 31, 533–538 (2013).

39. McInnes, L. & Healy, J. UMAP: Uniform Manifold Approximation and Projection for Dimension Reduction. ArXiv Mach. Learn. (2018).

40. Buchfink, B., Reuter, K. & Drost, H.-G. Sensitive protein alignments at tree-of-life scale using DIAMOND. Nat. Methods 18, 366–368 (2021).

41. Camaya, A. P. Stages of the symbiotic zooxanthellae–host cell division and the dynamic role of coral nucleus in the partitioning process: a novel observation elucidated by electron microscopy. Coral Reefs 39, 929–938 (2020).

42. Kempf, S. C. SYMBIOSIS BETWEEN THE ZOOXANTHELLA *SYMBIODINIUM* (= *GYMNODINIUM*) *MICROADRIATICUM* (FREUDENTHAL) AND FOUR SPECIES OF NUDIBRANCHS. Biol. Bull. 166, 110–126 (1984).

43. Lobo-da-Cunha, A., Alves, Â., Oliveira, E., Guimarães, F. & Calado, G. Endocytosis, lysosomes, calcium storage and other features of digestive-gland cells in cephalaspidean gastropods (Euopisthobranchia). J. Molluscan Stud. https://doi.org/10.1093/mollus/eyy034 (2018) doi:10.1093/mollus/eyy034.

44. Lobo-da-Cunha, A. Structure and function of the digestive system in molluscs. Cell Tissue Res. 377, 475–503 (2019).

45. Carmona, L., Pola, M., Gosliner, T. M. & Cervera, J. L. The end of a long controversy: systematics of the genus *Limenandra* (Mollusca: Nudibranchia: Aeolidiidae). Helgol. Mar. Res. 68, 37–48 (2014).

46. Kienberger, K. et al. *Aeolidia papillosa* (Linnaeus, 1761) (Mollusca: Heterobranchia: Nudibranchia), single species or a cryptic species complex? A morphological and molecular study. Zool. J. Linn. Soc. 177, 481–506 (2016).

47. Carmona, L., Cervera, J. L., Kumar, A. B. & Snehachandran, B. K. First record of the Aeolid *Anteaeolidiella fijensis* (Nudibranchia, Aeolidiidae) from India. Mar. Biodivers. 47, 823–830 (2017).

48. Carmona, L., Pola, M., Gosliner, T. M. & Cervera, J. L. Integrative taxonomy and biogeography of the genus *Bulbaeolidia* (Nudibranchia: Aeolidida). J. Molluscan Stud. 83, 440–450 (2017).

49. Karmeinski, D. et al. Transcriptomics provides a robust framework for the relationships of the major clades of cladobranch sea slugs (Mollusca, Gastropoda, Heterobranchia), but fails to resolve the position of the enigmatic genus Embletonia. BMC Ecol. Evol. 21, 226 (2021).

50. MolluscaBase Eds. MolluscaBase. Accessed through: World Register of Marine Species on 2025-06-30. VLIZ 10.14284/448 (2025).

51. Kempf, S. C. A ‘Primitive’ Symbiosis between the Aeolid Nudibranch Berghia Verrucicornis (A. Costa, 1867) and a Zooxanthella. J. Molluscan Stud. 57, 75–85 (1991).

52. Rudman, W. B. The taxonomy and biology of further aeolidacean and arminacean nudibranch molluscs with symbiotic zooxanthellae. Zool. J. Linn. Soc. 74, 147–196 (1982).

53. Cabrito, A., Galià-Camps, C. & Ballesteros, M. The risk of the nudibranch *Baeolidia rieae* (Mollusca: Gastropoda: Heterobranchia) in the international aquarium trade. https://doi.org/10.21411/CBM.A.92E8654F (2022) doi:10.21411/CBM.A.92E8654F.

54. Wägele, M. & Johnsen, G. Observations on the histology and photosynthetic performance of “solar-powered” opisthobranchs (Mollusca, Gastropoda, Opisthobranchia) containing symbiotic chloroplasts or zooxanthellae. Org. Divers. Evol. 1, 193–210 (2001).

55. McFarland, F. K. & Muller-Parker, G. Photosynthesis and Retention of Zooxanthellae and Zoochlorellae Within the Aeolid Nudibranch *Aeolidia papillosa*. Biol. Bull. 184, 223–229 (1993).

56. Burghardt, I. & Wägele, H. Interspecific differences in the efficiency and photosynthetic characteristics of the symbiosis of ‘solarpowered’ Nudibranchia (Mollusca: Gastropoda) with zooxanthellae. Rec. West. Aust. Mus. Suppl. 69, 1 (2006).

57. Goodheart, J. A. et al. Comparative morphology and evolution of the cnidosac in Cladobranchia (Gastropoda: Heterobranchia: Nudibranchia). Front. Zool. 15, 43 (2018).

58. Rudman, W. B. FURTHER STUDIES ON THE TAXONOMY AND BIOLOGY OF THE OCTOCORAL-FEEDING GENUS PHYLLODESMIUM EHRENBERG, 1831 (NUDIBRANCHIA: AEOLIDOIDEA). J. Molluscan Stud. 57, 167–203 (1991).

59. Quinlan, P. D. & Katz, P. S. State-dependent, visually guided behaviors in the nudibranch *Berghia stephanieae*. J. Exp. Biol. 226, jeb245213 (2023).

60. Borgstein, N. M., Burgués Palau, L., Parodi, B. A. & Laetz, E. M. J. Unraveling the *Phidiana* paradox: *Phidiana lynceus* can retain algal symbionts but its nocturnal tendencies prevent benefits from photosynthesis. Symbiosis 92, 245–255 (2024).

61. Ishikura, M. et al. Isolation of New *Symbiodinium* Strains from Tridacnid Giant Clam (*Tridacna crocea*) and Sea Slug (*Pteraeolidia ianthina*) Using Culture Medium Containing Giant Clam Tissue Homogenate. Mar. Biotechnol. 6, 378–385 (2004).

62. Loh, W., Cowlishaw, M. & Wilson, N. Diversity of *Symbiodinium* dinoflagellate symbionts from the Indo-Pacific sea slug *Pteraeolidia ianthina* (Gastropoda: Mollusca). Mar. Ecol. Prog. Ser. 320, 177–184 (2006).

63. FitzPatrick, S. K., et al. *Symbiodinium* diversity in the soft coral *Heteroxenia sp.* and its nudibranch predator *Phyllodesmium lizardensis*. Coral Reefs 31, 895–905 (2012).

64. Ziegler, M. et al. Thermal stress response in a dinoflagellate-bearing nudibranch and the octocoral on which it feeds. Coral Reefs 33, 1085–1099 (2014).

65. Yorifuji, M., Takeshima, H., Mabuchi, K., Watanabe, T. & Nishida, M. Comparison of *Symbiodinium* dinoflagellate flora in sea slug populations of the *Pteraeolidia ianthina* complex. Mar. Ecol. Prog. Ser. 521, 91–104 (2015).

66. Wecker, P. et al. Dinoflagellate diversity among nudibranchs and sponges from French Polynesia: Insights into associations and transfer. C. R. Biol. 338, 278–283 (2015).

67. Soon, N., Quek, Z. B. R., Pohl, S. & Wainwright, B. J. More than meets the eye: characterizing the cryptic species complex and Symbiodiniaceae communities in the reef-dwelling nudibranch *Pteraeolidia* ‘ *semperi* ’ (Nudibranchia: Aeolidioidea) from Singapore. J. Molluscan Stud. 89, eyad011 (2023).

68. Gafarova, E. et al. Gut Bacteriomes and Ecological Niche Divergence: An Example of Two Cryptic Gastropod Species. Biology 12, 1521 (2023).

69. Bernasconi, R., Stat, M., Koenders, A. & Huggett, M. J. Global Networks of *Symbiodinium*-Bacteria Within the Coral Holobiont. Microb. Ecol. 77, 794–807 (2019).

70. Pogoreutz, C. et al. Coral holobiont cues prime *Endozoicomonas* for a symbiotic lifestyle. ISME J. 16, 1883–1895 (2022).

71. Pogoreutz, C. & Ziegler, M. Frenemies on the reef? Resolving the coral–*Endozoicomonas* association. Trends Microbiol. 32, 422–434 (2024).

72. Wada, N. et al. High-resolution spatial and genomic characterization of coral-associated microbial aggregates in the coral *Stylophora pistillata*. Sci. Adv. 8, eabo2431 (2022).

73. Ng, M. S., Soon, N., Chang, Y. & Wainwright, B. J. Bacterial and Fungal Co-Occurrence in the Nudibranch, *Pteraeolidia semperi*. Life 12, 1988 (2022).

